# A genome-wide CRISPR knockout screen identifies host dependency factors and potential viral-induced cell death regulators for porcine deltacoronavirus infection

**DOI:** 10.64898/2026.10.06.755730

**Authors:** Thamonpan Laocharoensuk, Patricia A Boley, Rosemary Blake, Vayalena Drampa, Nicholas Parkinson, Finn Grey, Linda J Saif, Qiuhong Wang, Scott P Kenney

## Abstract

Porcine deltacoronavirus (PDCoV) is a globally circulating coronavirus of swine that causes severe enteric disease and mortality in neonatal piglets. Its broad host range and zoonotic potential raises concerns about continued spillover into human populations. However, the host factors that govern PDCoV infection remain largely unexplored. Identifying host factors involved in virus infection may uncover novel therapeutic targets for antiviral drug development or targets for gene editing to create resistant pigs. Here, we conducted a genome-wide CRISPR knockout (GeCKO) screen to identify host factors contributing to PDCoV infection in swine testicular (ST) cells. Small guide RNAs (sgRNAs) targeting the *ANPEP* gene, expressing the main receptor porcine aminopeptidase N for PDCoV, were enriched in PDCoV-resistant cells, demonstrating the validity of the screen. Moreover, we identified transmembrane protein 30A (TMEM30A), an accessory β-subunit of the P4-ATPase flippase complex, as a host dependency factor for PDCoV infection along with several potential PDCoV-induced cell death regulators, including DET1, IFNAR1, RFWD2, STAT1 and TP53. We further demonstrated that PDCoV attachment and internalization into cells were significantly impaired in TMEM30A knockout ST cells, suggesting that TMEM30A is important in the early stages of the PDCoV life cycle. Because TMEM30A also contributes to SARS-CoV-2 replication, future studies are warranted to investigate its mechanistic role in the context of coronavirus infection.

## 3. Introduction

Porcine deltacoronavirus (PDCoV), a member of the genus *Deltacoronavirus*, is a swine enteric coronavirus thought to have recently emerged from avian deltacoronaviruses. PDCoV causes mortality in neonatal piglets due to acute watery diarrhea, vomiting and severe dehydration (1,2). PDCoV was first identified in 2009 during a large-scale surveillance study in Hong Kong (3); however, its etiological role was not confirmed until 2014, when the first PDCoV-associated diarrhea outbreak occurred in the United States (1,4). Although PDCoV has shown a lower occurrence compared to other major swine pathogens, it continues to be present in US breeding herds (5) and in swine herds globally (6). Furthermore, PDCoV was detected in plasma samples from pediatric patients with acute febrile illness in Haiti (7), underscoring its zoonotic pathogen status (8). The persistence of viral circulation is concerning, as it provides ongoing opportunities for viral evolution, potentially leading to increased virulence or enhanced zoonotic potential.

PDCoV mainly utilizes aminopeptidase N (APN) as a receptor for attachment and entry (9,10), including on human cells (9,11). However, knockout of porcine APN does not completely block infection *in vitro* or *in vivo* (12–14), suggesting the presence of alternative viral entry pathways. Recently, several host factors associated with PDCoV infection have been identified using genome-wide CRISPR screens (15– 17). For example, the solute carrier family 35 member A1 (SLC35A1) is required for PDCoV adsorption by regulating sialic acid expression on the cell surface (15). C16orf62, a component of the multiprotein retriever complex, was shown to facilitate PDCoV binding and internalization through the regulation of cell surface APN (16). The heat shock protein 90 alpha family class B1 (HSP90AB1) was reported to promote PDCoV replication through its ATPase activity and prevent the proteasomal degradation of PDCoV nucleocapsid (N) protein (17). These findings underscore the need for further investigation into host factors associated with PDCoV infection to enhance our understanding of virus-host interaction, which could aid in antiviral development and genetic strategies such as selective breeding or gene modification for viral disease-resistant animals.

In this study, utilizing a genome-wide CRISPR knockout (GeCKO) screen, we identified multiple candidate host factors associated with PDCoV infection, including APN, along with host factors that potentially regulate PDCoV-induced cell death, including DET1 Partner Of COP1 E3 Ubiquitin Ligase (DET1), Interferon Alpha and Beta Receptor Subunit (IFNAR1), Ring Finger and WD Repeat Domain 2 (RFWD2), Signal Transducer and Activator of Transcription 1 (STAT1) and tumor protein 53 (TP53). Among the identified host factors, we further studied the role of TMEM30A, an accessory β-subunit of the P4-ATPase flippase complex, on PDCoV infection.

## 4. Materials and Methods

### 4.1 Plasmid construction

The lentiCas9-Blast (Addgene #52962) was used for generation of the ST and LLC-PK1 Cas9 cell lines (ST-Cas9 and LLC-PK1-Cas9). To construct the lentiviral sgRNA expression vector for TMEM30A knockout, the paired oligonucleotides (Table S1) of sgRNA were annealed and cloned into the linearized pKLV2-U6gRNA5(BbsI)-PGKpuro2ABFP-W (Addgene #67974) which was previously digested by BbsI restriction enzyme (R3539S, New England Biolabs).

### 4.2 Cells and viruses

Swine testicular (ST) cells, porcine kidney LLC-PK1, and human embryo kidney HEK293T cells were purchased from the American Type Culture Collection (ATCC). Aminopeptidase N knockout ST (APN KO ST) cells were a gift from Dr. Berend Jan Bosch (9). The PDCoV strain OH-FD22 passage 100 (P100) (1), was used in the survival screen, whereas passage 7 (P7) (GenBank accession No. MZ291567) (18) was used for validation experiments.

### 4.3 Lentiviral production and transduction

Lentivirus was produced as described previously (19) with some modifications. Briefly, co-transfection of the sgRNA expression plasmid or Cas9 expression plasmid, psPAX2 (Addgene #12260), and pMD2.G (Addgene #12259) was performed using Lipofectamine™ LTX reagent with PLUS reagent (Thermo Fisher). The supernatants collected at 24 and 48 hours post transfection were pooled, filtered through sterile 0.22 µm PES syringe filters (Millipore), aliquoted and stored at -80°C. Transduction was performed in the presence of 8 µg/mL polybrene (Santa Cruz Biotechnology).

### 4.4 Generation of ST and LLC-PK1 cells stably expressing Cas9

ST and LLC-PK1 cells were transduced with Cas9-Blast lentivirus. At 48 hours post transduction, the transduced cells were subjected to blasticidin (Research Products International) selection (10 µg/mL). Clonal cells were isolated by limiting dilution and Cas9 expression in the resulting clones was confirmed by Western blot (Fig. S1).

### 4.5 Western blotting

Cell lysates were prepared using 300 µL of RIPA lysis buffer (Thermo Fisher). Equal amounts of protein were resolved by SDS-PAGE and transferred onto PVDF membranes (Bio-Rad). Blotted membranes were blocked with blocking buffer (Li-COR) for 2 hours at room temperature, followed by overnight incubation at 4 °C with a primary antibody against *Streptococcus pyogenes* (Sp) Cas9 (A01935-40, GenScript). After washing, membranes were incubated with an AlexaFluor 488-conjugated secondary antibody (A11029, Thermo Fisher) for 2 hours at room temperature. Visualization was carried out using the iBright FL1500 Imaging System (Thermo Fisher).

### 4.6 Genome-wide CRISPR knockout (GeCKO) screening and data analysis

A genome-wide porcine sgRNA library consisting of 80,510 guides targeting 20,517 genes (Table S2) was developed by the groups led by Dr. Finn Grey and Dr. Nicholas Parkinson, based on the annotation of the pig genome version 11.1 (Sscrofa11.1). The pig GeCKO cell libraries were generated by lentiviral transduction of the ST-Cas9 cells at a multiplicity of infection (MOI) of 0.3 (20). Transduced cells were subjected to puromycin (Invivogen) selection (2 µg/mL) at 48 hours post transduction. At 6 days post transduction, the porcine GeCKO cells were harvested for genomic DNA (gDNA) extraction using the Quick-DNA Midiprep Plus Kit (Zymo Research) and subjected to Illumina sequencing (NextSeq1000). The uninfected library-transduced cells served as negative control and were used to determine library coverage. For the survival screen, the GeCKO cell library was challenged with PDCoV (MOI = 0.1) for 3 consecutive rounds, referred to as the first, second, and third selections. On 2 days post-inoculation (dpi), PDCoV-resistant cells were harvested and expanded for deep sequencing and the next round of virus challenge. gDNA was extracted from the surviving cells from each selection and subjected to deep sequencing. The sequencing data was analyzed using MAGeCK Robust Rank Aggregation (RRA) method (21) on the Galaxy web app (22). The top ten candidate host factors from each selection were combined into a total of 13 unique genes and uploaded to ShinyGO (version 0.85.1) for pathway enrichment analysis based on the Kyoto Encyclopedia of Genes and Genomes (KEGG) pathway database. The following parameters were used for the analysis: false discovery rate (FDR) < 0.05, with the minimum genes per pathway set to 2 to accommodate the small number of top candidate genes in the list.

### 4.7 Generation of TMEM30A knockout (KO) cell clones

ST-Cas9 and LLC-PK-Cas9 cells were transduced with lentivirus expressing sgRNAs targeting exons 3, 4 and 5 of TMEM30A. Following puromycin selection, single cell clones were isolated by limiting dilution. INDELs were confirmed by Sanger sequencing, and editing efficiency was assessed using the Tracking of Indels by Decomposition (TIDE) program (23). Primers targeting each exon are listed in Table S3.

### 4.8 MTS proliferation assay

Cells were seeded into 96-well plates at a density of 5,000 cells per well. At 48 hours post seeding, 20 µL of CellTiter 96® AQueous One Solution reagent (Promega) was added to each well. The plate was incubated for 2 hours at 37°C, and the optical density (OD) was measured at 450 nm using an ELISA plate reader (FilterMax F5, Molecular Devices). Each group included five replicates to ensure the reliability of the results and the experiments were repeated twice.

### 4.9 RT-qPCR

Cells were infected with PDCoV in the presence of trypsin (5 µg/mL) (Gibco) or 1% pancreatin (Sigma). Total RNA was extracted from the supernatants harvested at 24 hours post-inoculation (hpi) using TRIzol^®^ Reagent (Thermo Fisher) and subsequently treated with DNase I (Thermo Fisher) to remove residual gDNA. A one-step RT-qPCR was performed using the TaqMan™ Fast Virus 1-Step Master Mix (Thermo Fisher). Primers and probe targeting the PDCoV nucleocapsid (N) gene were derived from Chen et al. (24). Viral genomic copy numbers were determined by extrapolation from a standard curve generated using a plasmid carrying the PDCoV N gene generated in our lab.

### 4.10 Infectious virus titration

The quantification of virus titer from the cell culture supernatant harvested at 12 and 24 hpi was performed by determining median tissue culture infectious dose (TCID_50_) using LLC-PK1 cells as previously described (1).

### 4.11 Binding and internalization assays

To determine PDCoV binding, wildtype (WT) and TMEM30A KO ST cells were seeded and incubated with PDCoV at MOI of 5 at 4°C to allow virus binding to cell surface. The inocula were then removed after 1 hour and cells were washed thrice with ice-cold PBS (pH 7.4). For the internalization assay, cells were incubated with PDCoV at MOI of 1 at 4°C for 1 hour. Following removal of unbound viral particles, cells were incubated at 37°C for 1 hour to allow for internalization. The non-internalized virus particles were then removed by washing with ice-cold acidic PBS (pH 1.3) three times. Finally, cells from both assays were lysed with TRIzol^®^ Reagent and the cell lysates were subjected to RNA extraction and RT-qPCR.

### 4.12 Statistical analysis

Statistical analyses were performed using GraphPad Prism version 10.2.3. Data is presented as means ± standard deviation (SD). A p-value of less than 0.05 is considered statistically significant. The employed statistical tests are indicated in the corresponding figure legend.

## 5. Results

### 5.1 Genome-wide CRISPR knockout screen identifies candidate host dependency factors and potential virus-induced cell death regulators for PDCoV infection in ST cells

To identify host factors essential for cell survival in response to PDCoV infection, we performed a genome-wide CRISPR knockout screen in ST cells. The schematic diagram of the screening workflow is illustrated in Fig. 1A. The sgRNA abundance in the uninfected library and PDCoV-resistant cells from each selection was determined by Illumina sequencing. In total, 95.94% (77229/80510) of the originally designed sgRNA sequences were retained in the ST GeCKO cell collection (Fig. 1B).

**Fig. 1.**
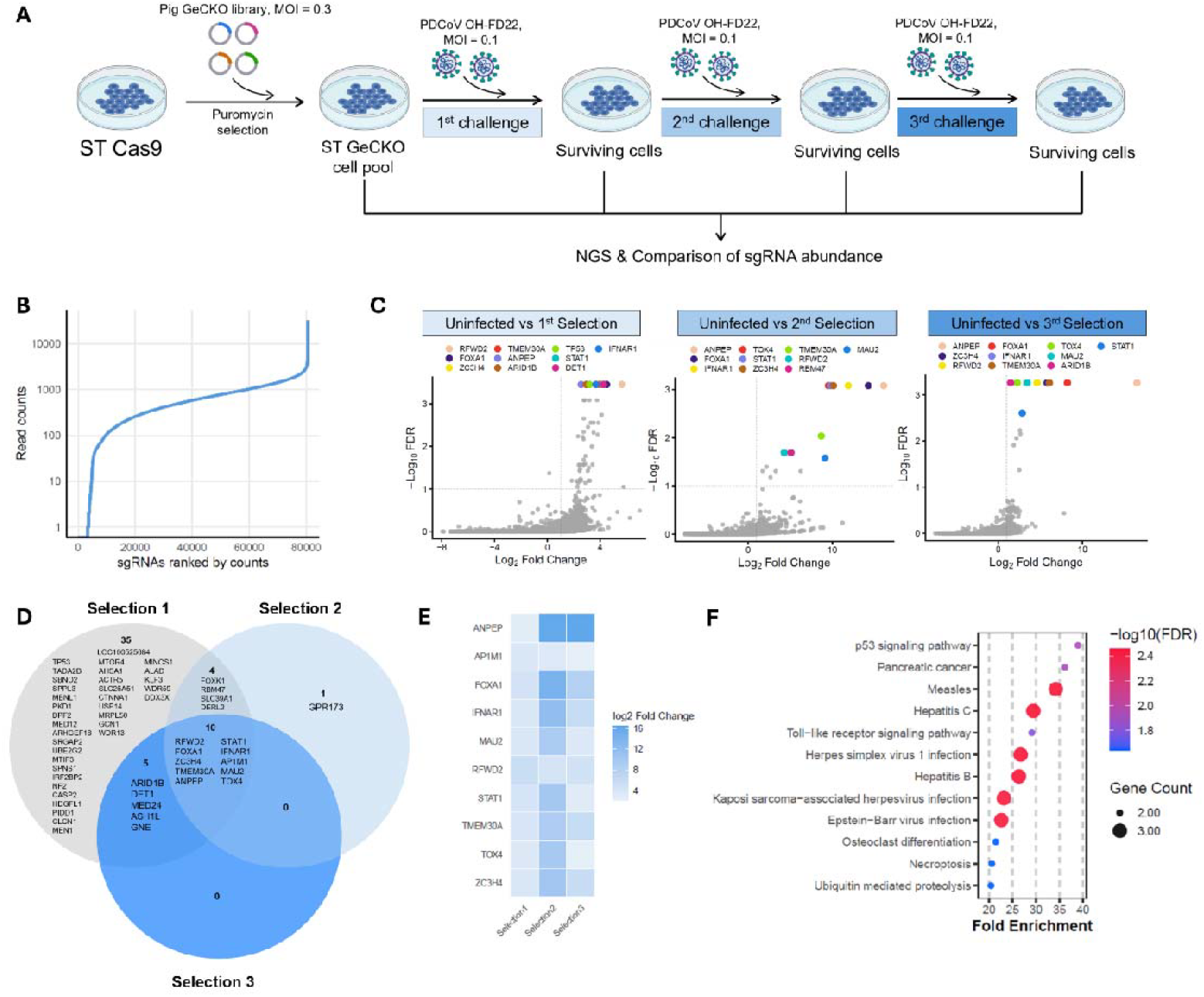
GeCKO screen identified potential candidate host factors associated with PDCoV infection in ST cells. (A) Schematic diagram of GeCKO screening workflow created using BioRender (B) The library coverage was determined by deep sequencing the uninfected GeCKO library-transduced population. The curve indicates the sgRNA distribution. (C) Volcano plots showing the top 10 most enriched genes from three selections analyzed by MAGeCK test tool (Galaxy Version 0.5.9.2.1). Dashed lines indicate the cutoff for significant candidate host genes (FDR <0.1 and log2 fold change ≥1). (D) Venn diagram showing significant genes (FDR <0.1 and log2 fold change ≥1) from three challenges. (E) Heatmap comparing log2 fold changes of ten significant overlapping genes identified across three selections, with gene names ordered alphabetically. (F) Dot plot showing the top enriched KEGG pathways represented by the top candidate genes. The details about genes representing the enriched pathways can be found in Table S4.

We employed the MAGeCK test tool to compare significantly enriched sgRNAs present in surviving cells from each selection to those from an uninfected GeCKO library transduced cell population. A highly enriched sgRNA indicates that the knockout of the targeted gene confers a resistance to PDCoV-induced cell death. MAGeCK analyses revealed the *ANPEP* gene, encoding porcine aminopeptidase N (pAPN), a known primary receptor for PDCoV, was selected in all three selections and as the top candidate based on the robust rank aggregation (RRA) score in the 2^nd^ and 3^rd^ selections (Fig. 1C), demonstrating the effectiveness and validity of our CRISPR knockout screen.

The top 10 most positively selected genes were consistent, with 7/10 being found in all three rounds of selection, including *ANPEP, RFWD2* (*COP1*), Forkhead box A1 (*FOXA1*), zinc finger CCCH-type containing 4 (*ZC3H4*), *TMEM30A* (*CDC50A*), *STAT1*, and *IFNAR1* (Fig. 1C). There were ten significant candidate host factors that were shared among three selections (Fig. 1D and 1E). Adaptor-related protein complex AP-1 mu subunit 1 (*AP1M1*) was not initially in the list of top ten candidate genes based on RRA score but was considered significant according to the cutoff criteria (false discovery rate; FDR < 0.1 and log2 fold change ≥ 1). Pathway enrichment analysis of the top candidate genes from three selections (13 genes in total) based on the KEGG database showed enriched pathways related to not only various viral infections, but also regulation of cell death pathways such as p53 signaling pathway, necroptosis, and ubiquitin-mediated proteolysis (Fig. 1F and Table S4).

### 5.2 TMEM30A is a host dependency factor for PDCoV infection in swine cell lines

Because TMEM30A was previously implicated in SARS-CoV-2 infection (25), we focused on its impact on PDCoV replication. TMEM30A is the β-subunit for many P4-ATP flippases (26) and localizes to the cell membrane and membrane organelles (Table 1). Clonal TMEM30A KO cell lines were generated using a CRISPR/Cas9 system. Sanger sequencing confirmed that these clonal cell lines carried a one nucleotide insertion predicted to induce a frameshift mutation that effectively disrupts TMEM30A coding, resulting in the expression of the N-terminal 130 amino acid residues of a natural TMEM30A followed by seven extra amino acids (131SLCEISR137) due to the frame shift (Fig. 2A-C and Fig. S2). We observed no negative impact of TMEM30A knockout on normal cell proliferation or morphology (Fig. 2D and 2E). However, a significant decrease in viral titers and PDCoV N gene copy numbers post-infection was observed in the TMEM30A KO ST cells compared with parental ST cells (Fig. 2F and 2G). Similar findings were also observed in TMEM30A KO LLC-PK1 cells (Fig. 2H and 2I). Together, these results suggest that TMEM30A promotes PDCoV infection in both ST and LLC-PK1 cell lines.

**Table 1.**
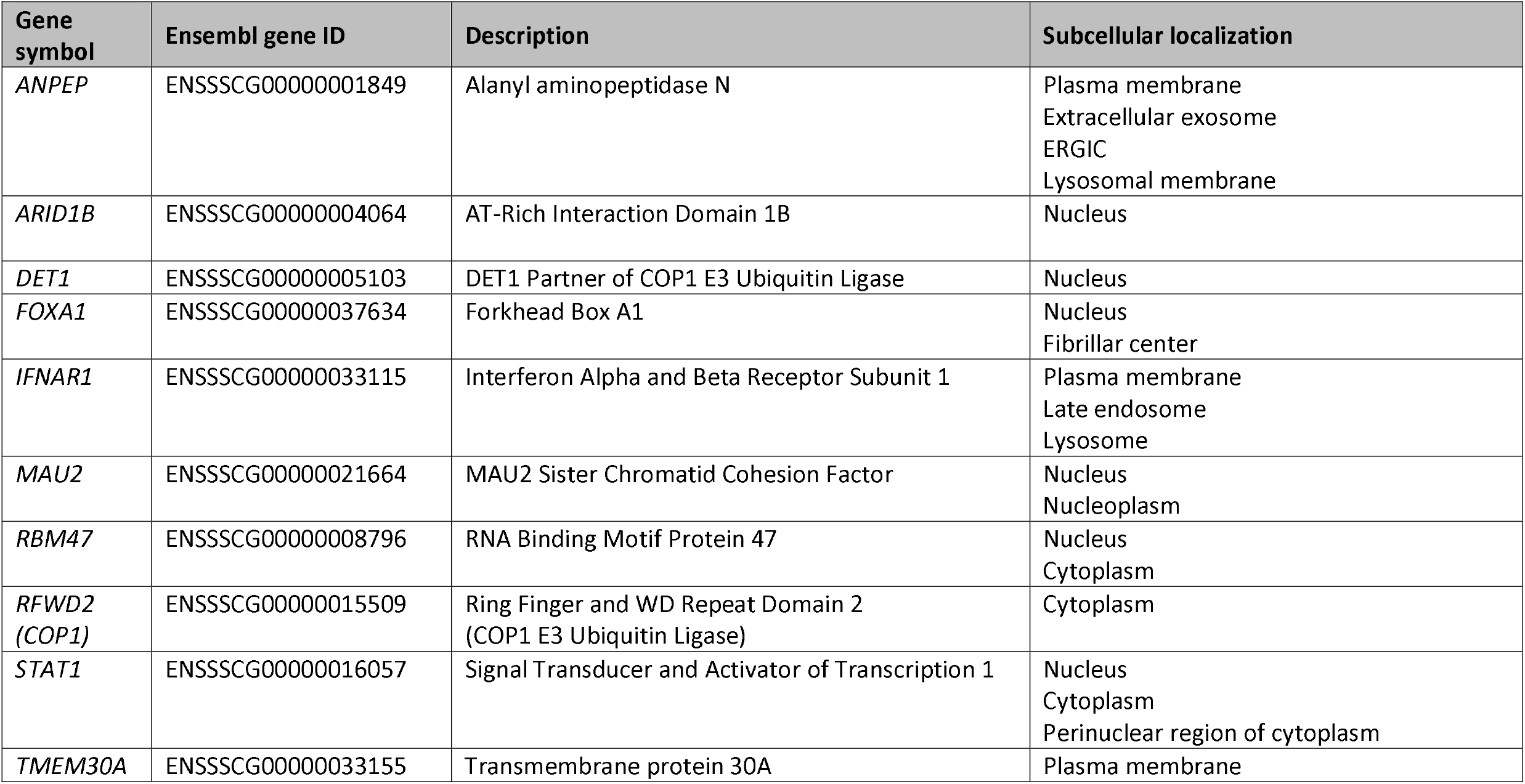
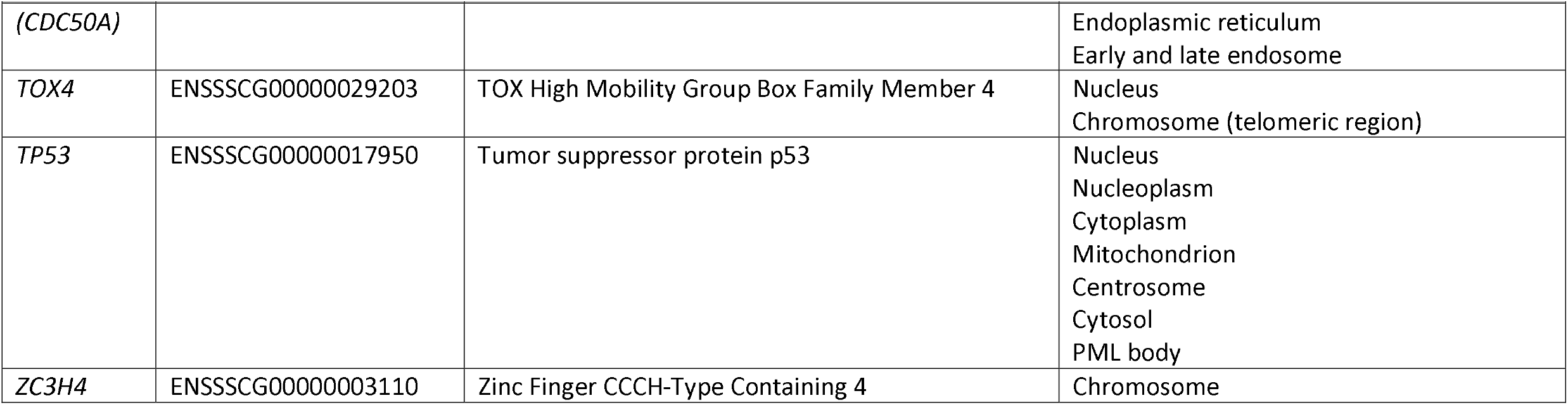
Top candidate host factors identified from GeCKO screen in ST cells. The subcellular localizations for each gene are based on UniProt annotation. Genes are ordered alphabetically.

**Fig. 2.**
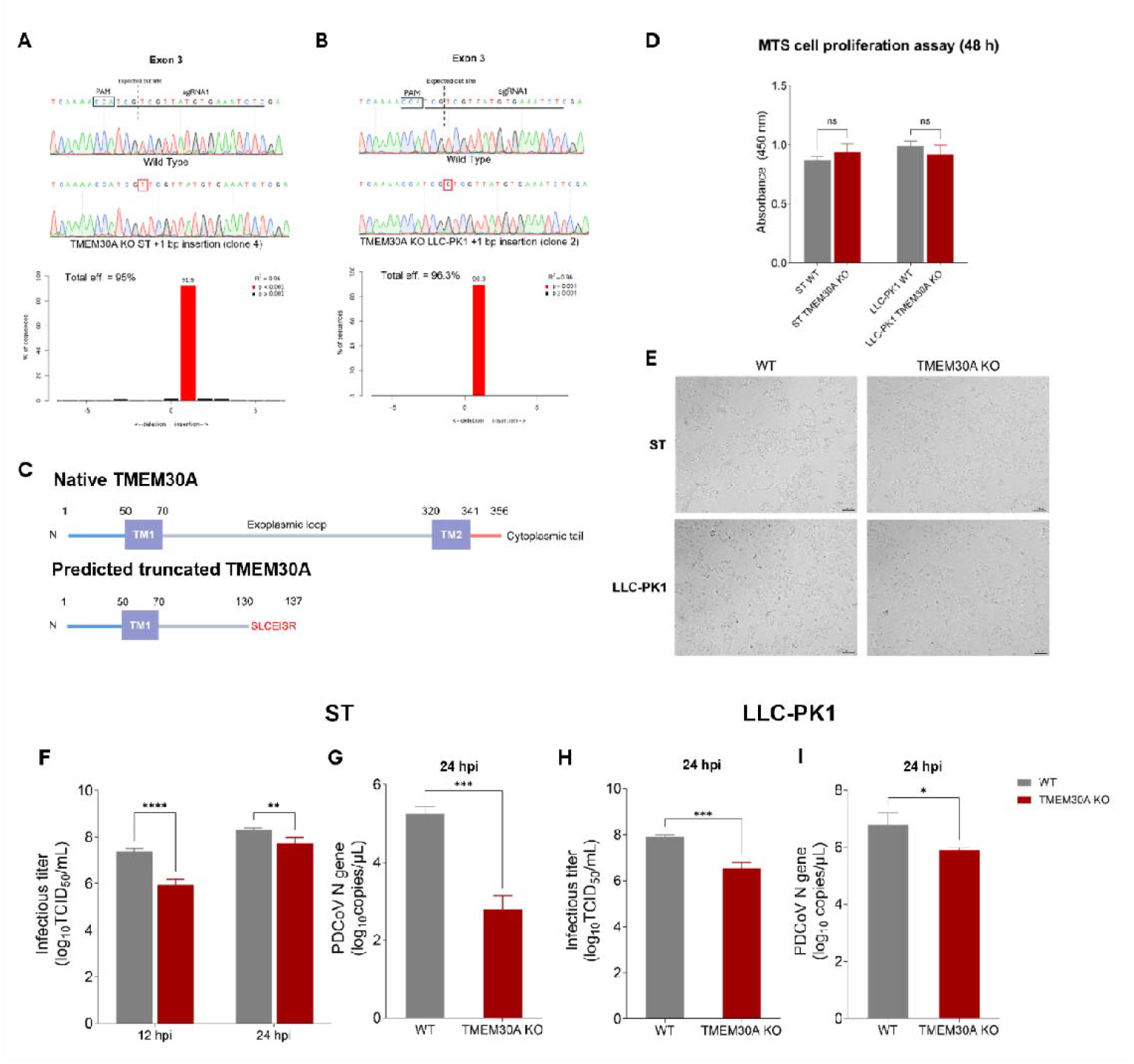
TMEM30A contributes to PDCoV infection in swine cell lines. (A) and (B) Sanger sequencing confirming a single nucleotide insertion (red boxes) in exon 3 of the TMEM30A gene in both ST and LLC-PK1 cell lines. The sgRNA binding site, expected cut site and PAM sequences are highlighted. TIDE analyses showed that the primary introduced mutations were a single insertion with high editing efficacy. (C) Schematic diagram of native and predicted truncated TMEM30A. The insertion mutations result in earlier termination of TMEM30A expression at amino acid residue 130, followed by seven amino acids (SLCEISR). (D) Evaluation of TMEM30A disruption on cell proliferation by MTS proliferation assay (One-way ANOVA followed by Šídák multiple comparisons test, data are from one experiment representative of 2 independent trials). (E) Microscopic morphology of TMEM30A KO and WT cells (100x magnification, scale bar = 100 µm). (F) to (I) Effects of TMEM30A knockout on PDCoV infection in ST and LLC-PK1 cells (Two-way ANOVA followed by Šídák’s multiple comparisons test for Fig. 2F and Unpaired, Two-tailed Student’s t-test for Fig. 2G-2I, n = 3 replicates). ns: non-significant; *: p < 0.05; **: p < 0.01 ***: p < 0.001; ****: p < 0.0001.

### 5.3 TMEM30A contributes to the early steps of PDCoV life cycle

After we confirmed the involvement of TMEM30A in PDCoV infection, we further investigated which stages in the PDCoV life cycle were affected by TMEM30A disruption. To do this, PDCoV binding and internalization assays were performed. We found that PDCoV showed significant reduction in binding to TMEM30A KO ST cells compared with wildtype ST cells (Fig. 3A). APN KO ST cells were used as a control. We observed comparable reduction in PDCoV N gene copies in APN KO and TMEM30A KO cells. PDCoV internalization was also reduced in the internalization assay (Fig. 3B). Although infectious viral titers were significantly reduced in both the cell lysate and supernatant of KO cells compared to WT cells (Fig. 3D), the ratio of extracellular to cell-associated infectious virus did not differ significantly (Fig. 3C), suggesting that TMEM30A disruption did not alter viral release. Collectively, these findings suggest that TMEM30A plays a role in entry steps of PDCoV infection rather than in replication or release.

**Fig. 3.**
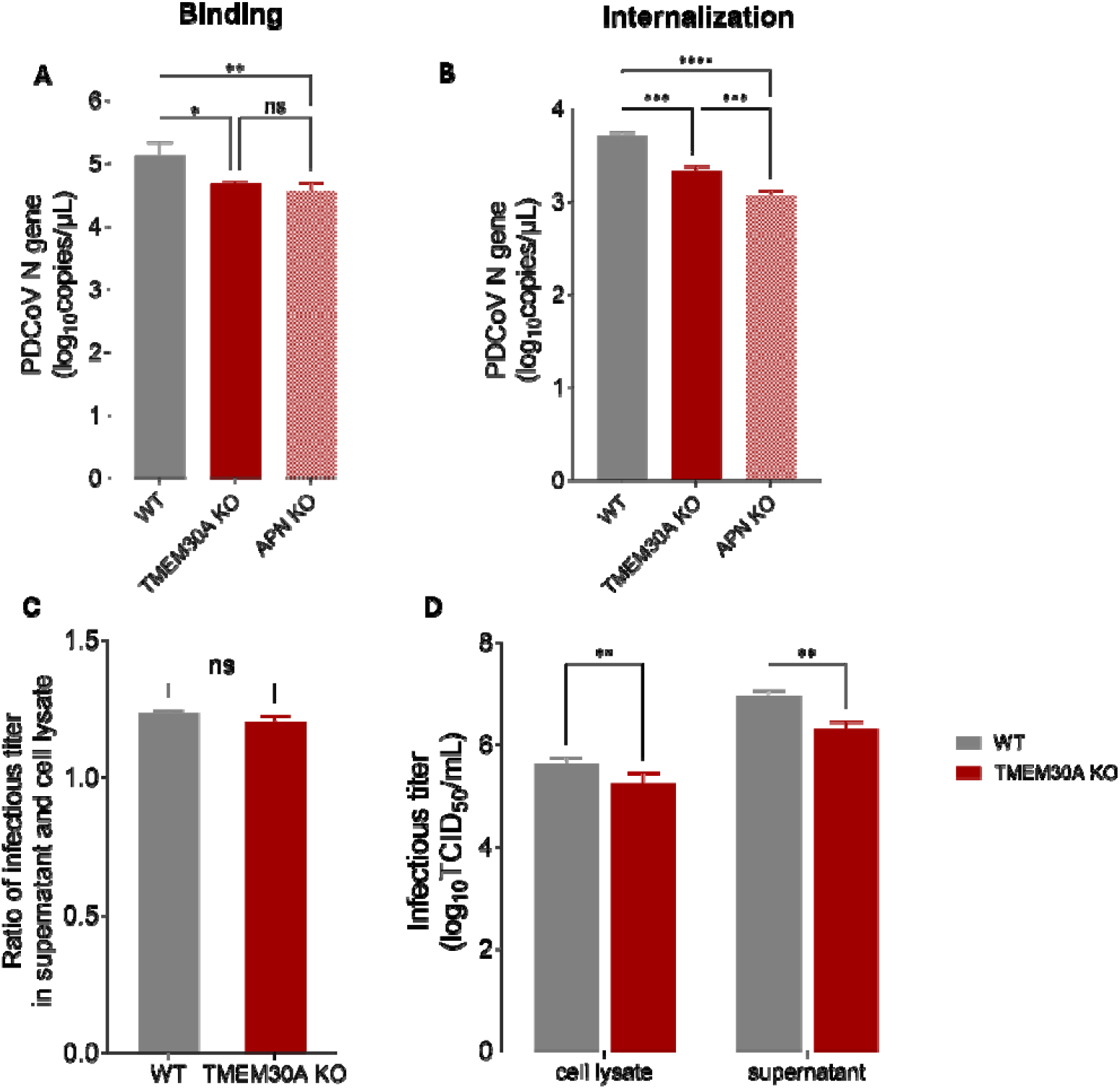
TMEM30A contributes to the early steps in PDCoV life cycle in ST cells. (A) Effects of TMEM30A disruption in PDCoV attachment (A) and internalization (B). APN KO cells were used as a control (One-way ANOVA followed by Tukey’s multiple comparisons test, n = 3 independent experiments). (C) and (D) Cells were infected with PDCoV at MOI of 0.5, cell lysates and supernatant were then harvested at 12 hpi for infectious virus titration (Unpaired, two-tailed Student’s t-test for Fig. 3C and Two-way ANOVA followed by Šídák’s multiple comparisons test for Fig. 3D). ns: non-significant; *: p < 0.05; **: p < 0.01 ***: p < 0.001; ****: p < 0.0001.

## 6. Discussion

Coronaviruses pose a significant global threat to animal and human health. The ability of PDCoV to cross species barriers (9,27,28), infect humans (7,8), and to continuously circulate in swine breeding herds (6) has raised concerns about its potential to evolve to become a more efficient zoonotic pathogen. This highlights an urgent need to identify novel therapeutic targets to effectively combat PDCoV infection.

In this study, we performed a GeCKO screen and identified multiple candidate host factors involved in PDCoV infection in swine-derived cells. As expected, *ANPEP* gene, encoding the primary entry receptor aminopeptidase N (APN, also known as CD13) for PDCoV, was identified as the top enriched gene during selection, confirming the validity of our screening strategy. One round of selection appeared to be effective at identifying candidate genes with the top 7 enriched genes being conserved across all three selection events (Figure 1C). Previous genome-wide CRISPR knockout screens for PDCoV have used porcine LLC-PK1 (15,17) and human Huh7 cells (16) to identify host factors essential for PDCoV infection. TMEM30A and the other top 10 candidate host factors identified in our screens, with the exception for APN, were not reported among the top candidate host factors found in the previous studies. These differences may reflect variations in cell type, viral strain, library design, selection conditions, or analysis methodology (Table 2).

**Table 2.**
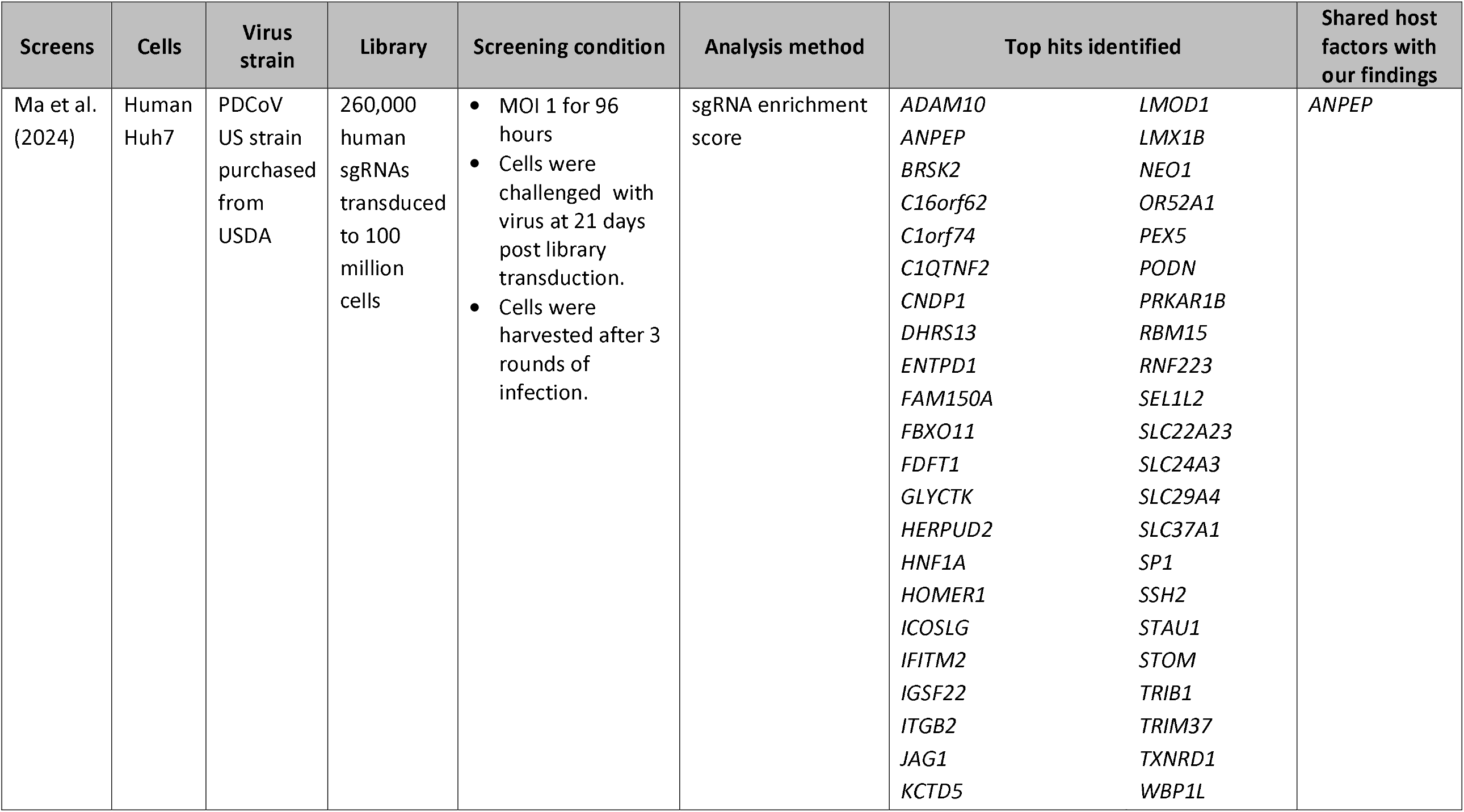
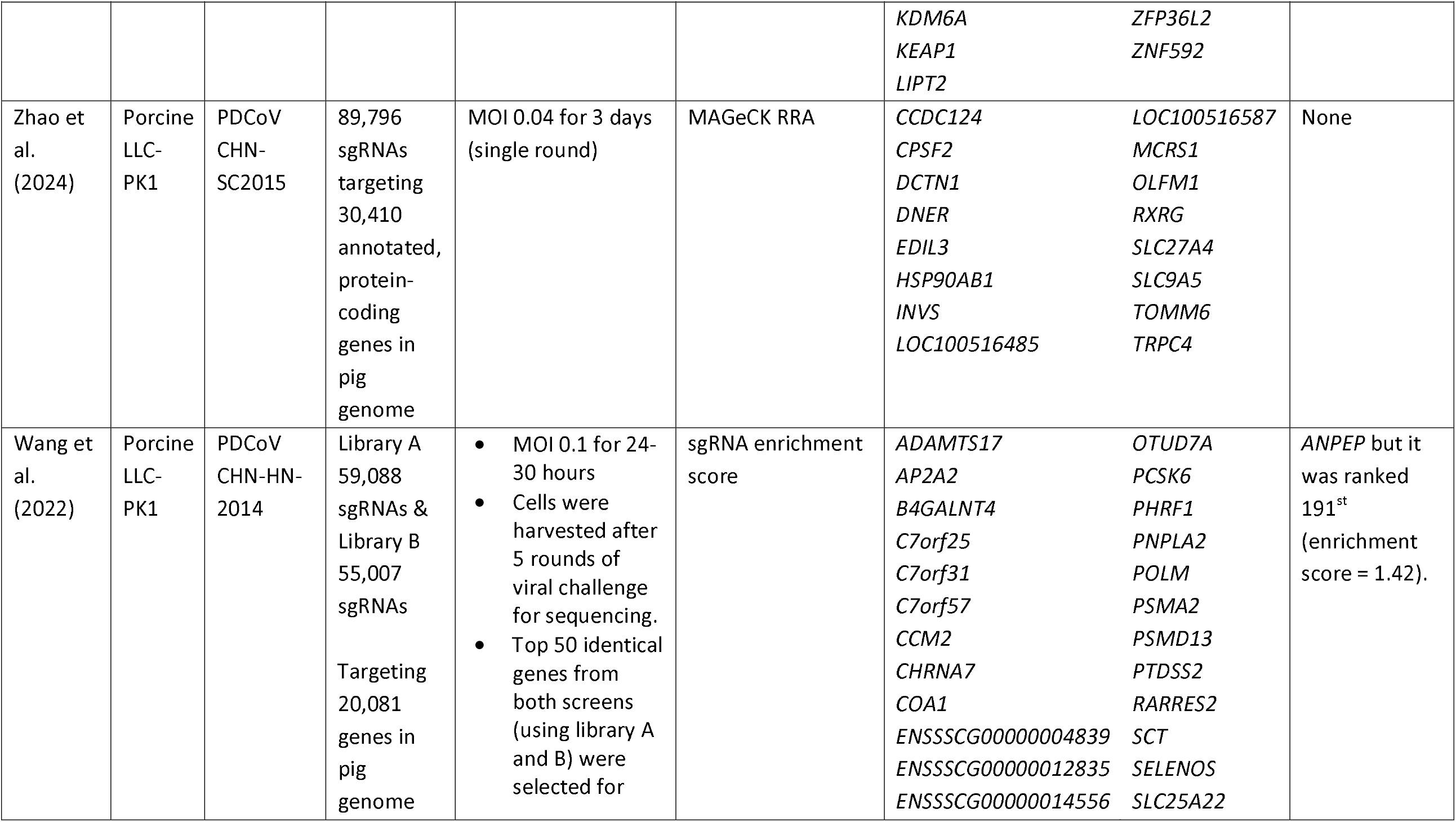
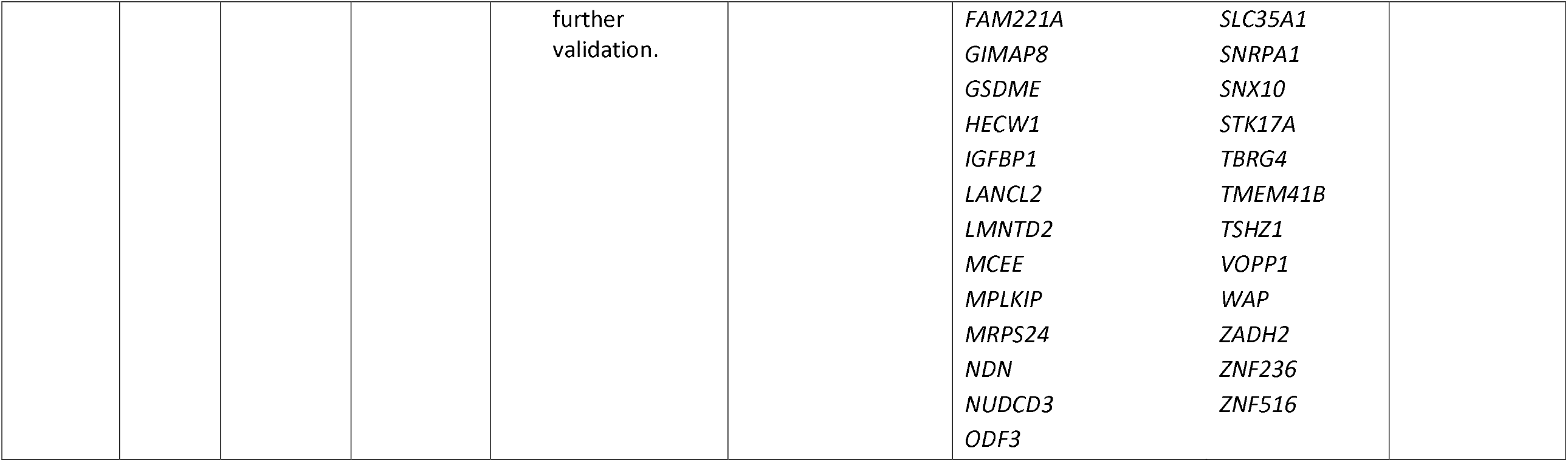
Comparative analysis of screening strategies and sgRNA/gene hits identified from previously published CRISPR KO screens against PDCoV.

In addition to candidate host dependency factors, our screen uncovered potential regulators of virus-induced cell death, including DET1, IFNAR1, RFWD2, STAT1, and TP53. As expected, the pathways represented by these genes are associated with regulation of cell death, including p53 signaling, necroptosis, and ubiquitin-mediated proteolysis (Figure 1F). *TP53* encodes a tumor suppressor protein p53, which has also been implicated in antiviral immunity in addition to its physiological roles in cell cycle regulation and programmed cell death in response to stress or injury (29,30). p53 is negatively regulated by ubiquitin proteasome degradation mediated by several E3 ubiquitin ligases, including COP1 (encoded by *RFWD2*) and its interacting partner DET1, both identified in our screen. p53 is also influenced by type I IFN (IFN-α/β) signaling (29) and has been established as a host restriction factor for several coronaviruses, including swine enteric coronaviral pathogens (31,32). Previous studies have shown a complex role of p53 in porcine epidemic diarrhea virus (PEDV), another swine enteric coronaviral pathogen. PEDV nucleocapsid (N) protein can interact with p53 to promote cell cycle arrest at S-phase, providing a favorable environment for early viral replication (33). Later in infection, PEDV N instead induces COP1-mediated p53 degradation to suppress apoptosis and facilitate viral dissemination (34). Although *TP53* can be enriched in survival screens without viral challenge, its enrichment in our screen may be related to viral selection. However, the effect of general cell fitness cannot be excluded as an unchallenged library control harvested at the same time was not utilized. Therefore, it would be interesting to investigate whether PDCoV activates COP1-mediated p53 degradation in a manner similar to PEDV.

The type I IFNs play a key bifunctional role in host defense against viral infection by conferring an antiviral state in uninfected cells and promoting apoptosis in virally infected cells (35). Interestingly, we observed that sgRNAs targeting IFNAR1 and its downstream mediator, STAT1, were enriched across all three selections. However, Jung et al. showed that PDCoV can induce apoptosis in ST and LLC-PK1 cells (36). This suggests that, in this context, knockout of IFNAR1 and STAT1 may inhibit PDCoV-induced cell death through apoptosis in infected cells, which could explain the enrichment of these two host factors in cells that survived PDCoV challenge.

Among the factors identified, we prioritized TMEM30A for further validation studies, as TMEM30A has been implicated in SARS-CoV-2 entry (25). Our findings showed that TMEM30A also contributes to PDCoV infection. TMEM30A, also known as CDC50A, is an essential accessory β-subunit of P4-ATPase flippase complex, which helps maintain the asymmetric distribution of phospholipids across cell membranes (26). While up to 14 types of P4-ATPases are present in mammalian cells, posing challenges for studying their physiological functions due to functional redundancy and overlapping tissue distribution, the TMEM30 accessory subunit family consists of only three members, with TMEM30A being the most predominant (26). Physiologically, TMEM30A is essential for myoblast fusion and proliferation and migration of satellite cells, two key processes involved in skeletal muscle regeneration following injury (37). TMEM30A is also required for vesicle-mediated protein trafficking, particularly in normal insulin maturation and glucose-induced insulin secretion in pancreatic β cells (38). Besides SARS-CoV-2, TMEM30A was identified as a host entry factor for murine norovirus (MNV) (39), although its specific role was not elucidated until recently, when TMEM30A was shown to facilitate MNV binding by maintaining lipid order and membrane fluidity (40). TMEM30A was also reported as an alternative entry receptor for SARS-CoV-2 by interacting with the N-terminal domain of SARS-CoV-2 spike protein, facilitating viral-host membrane fusion, potentially in tissues where the canonical entry factor ACE2 expression was low (25). However, the detailed mechanisms have not been elucidated. It is notable that PDCoV, like SARS-CoV-2, belongs to the family *Coronaviridae* and both can jump across species barriers producing infection in multiple hosts. Identifying host factors conserved across viruses within the same family could aid in the discovery of therapeutic targets for pan-antiviral development.

In summary, our study identified TMEM30A as a host dependency factor for PDCoV infection. The knockout of TMEM30A led to a significant reduction in both viral genomic copy numbers and infectious viral titers. We further showed that TMEM30A likely plays a role in early steps in the PDCoV life cycle. Future studies are warranted to elucidate its molecular mechanism in the context of PDCoV infection.

## 8. Author statements

### 8.1 Author contributions

**TL:** Conceptualization, Formal analysis, Investigation, Methodology, Visualization, Writing – original draft. **PAB:** Methodology, Investigation, Writing – review and editing. **LJS:** Resources, Writing – review and editing. **RB:** Methodology, Writing – review and editing. **VD:** Methodology, Writing – review and editing. **NP:** Formal analysis, Methodology, Writing – review and editing. **FG:** Funding acquisition, Resources, Methodology, Writing – review and editing. **QW:** Resources, Supervision, Writing – review and editing. **SPK:** Conceptualization, Funding acquisition, Methodology, Investigation, Supervision, Writing – review and editing.

## 8.4 Acknowledgements

We thank Dr. Wirat Pipatpongpinyo from Research and Analytical Service Core (RASC) for technical assistance with Illumina Nextseq sequencing and Kenney lab members for their valuable feedback and insights. We also thank Dr. Berend J Bosch from Utrecht University for kindly providing APN KO ST cells.

