## Supplementary Data for "A genome-wide CRISPR knockout screen identifies host dependency factors and potential viral-induced cell death regulators for porcine deltacoronavirus infection"

**Table S1** sgRNA sequences for generation of lentiviral sgRNA expression vector. Underlined sequences are 5′ overhangs for restriction enzyme BbsI used for cloning.

| Name | Sequence (5′ to 3′) |
|---|---|
| TMEM30A_guide1_F | CACCGAGATTTCACATAACGACGA |
| TMEM30A_guide1_R | AAACTCGTCGTTATGTGAAATCTC |
| TMEM30A_guide2_F | CACCTCTAGTGGAATGTCTCCAGA |
| TMEM30A_guide2_R | AAACTCTGGAGACATTCCACTAGA |
| TMEM30A_guide3_F | CACCGACAAACCAATTGCTCCTTG |
| TMEM30A_guide3_R | AAACCAAGGAGCAATTGGTTTGTC |

**Table S2** Details about the porcine GeCKO library and genes targeted by guide RNAs

| Genes | Count |
|---|---|
| Targeted protein-coding genes | 20,200 |
| Targeted miRNA genes | 317 |
| Genes with 1 guide | 393 |
| Genes with 2 guides | 292 |
| Genes with 3 guides | 309 |
| Genes with 4 guides | 19,523 |
| **Guide RNAs** | **Count** |
| Targeting guides | 79,993 |
| Non-targeting guides | 517 |
| Total guides | 80,510 |

**Table S3** Primers used for TMEM30A knockout validation

| Name | Sequence (5′ to 3′) | PCR product size (bp) |
|---|---|---|
| TMEM30A_exon3_F | AGAAATCACCCCAGTTCCTGTT | 555 |
| TMEM30A_exon3_R | CCCAGAATGCTCTAGACTCATCAA | |
| TMEM30A_exon4_F | AATGCAGCTGTGTTAAGGGATG | 599 |
| TMEM30A_exon4_R | GTTGGTGGTAAAGGTAGTCCA | |
| TMEM30A_exon5_F | TCAGCTAGCCATGTAGGGAA | 506 |
| TMEM30A_exon5_R | TGGCATAAAGTTAAGCAAAGTTGT | |

**Table S4** Top enriched pathways that the top candidate genes represented based on KEGG pathway database

| Pathway | Genes | FDR | Fold Enrichment |
|---|---|---|---|
| Path:ssc04115 p53 signaling pathway | COP1 TP53 | 0.012059 | 38.93491 |
| Path:ssc05212 Pancreatic cancer | STAT1 TP53 | 0.012059 | 36.15385 |

| | | | |
|---|---|---|---|
| Path:ssc05162 Measles | STAT1 IFNAR1 TP53 | 0.00348 | 34.19958 |
| Path:ssc05160 Hepatitis C | STAT1 IFNAR1 TP53 | 0.00348 | 29.42755 |
| Path:ssc04620 Toll-like receptor signaling pathway | STAT1 IFNAR1 | 0.016405 | 29.0893 |
| Path:ssc05168 Herpes simplex virus 1 infection | STAT1 IFNAR1 TP53 | 0.00348 | 26.73348 |
| Path:ssc05161 Hepatitis B | STAT1 IFNAR1 TP53 | 0.00348 | 26.36218 |
| Path:ssc05167 Kaposi sarcoma-associated herpesvirus infection | STAT1 IFNAR1 TP53 | 0.003654 | 23.14728 |
| Path:ssc05169 Epstein-Barr virus infection | STAT1 IFNAR1 TP53 | 0.003654 | 22.59615 |
| Path:ssc04380 Osteoclast differentiation | STAT1 IFNAR1 | 0.022908 | 21.4472 |
| Path:ssc04217 Necroptosis | STAT1 IFNAR1 | 0.022908 | 20.57536 |
| Path:ssc04120 Ubiquitin mediated proteolysis | COP1 DET1 | 0.022908 | 20.40943 |

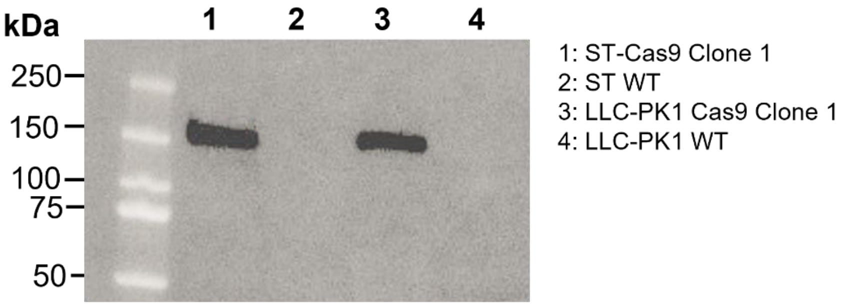


**Fig. S1** Confirmation of Cas9 expression by Western blot. Cell lysates from clonal Cas9 and parental cells were analyzed by Western blot using a specific antibody against *Streptococcus pyogenes (Sp)* Cas9. Samples were loaded along with a prestained protein ladder (Bio-Rad).

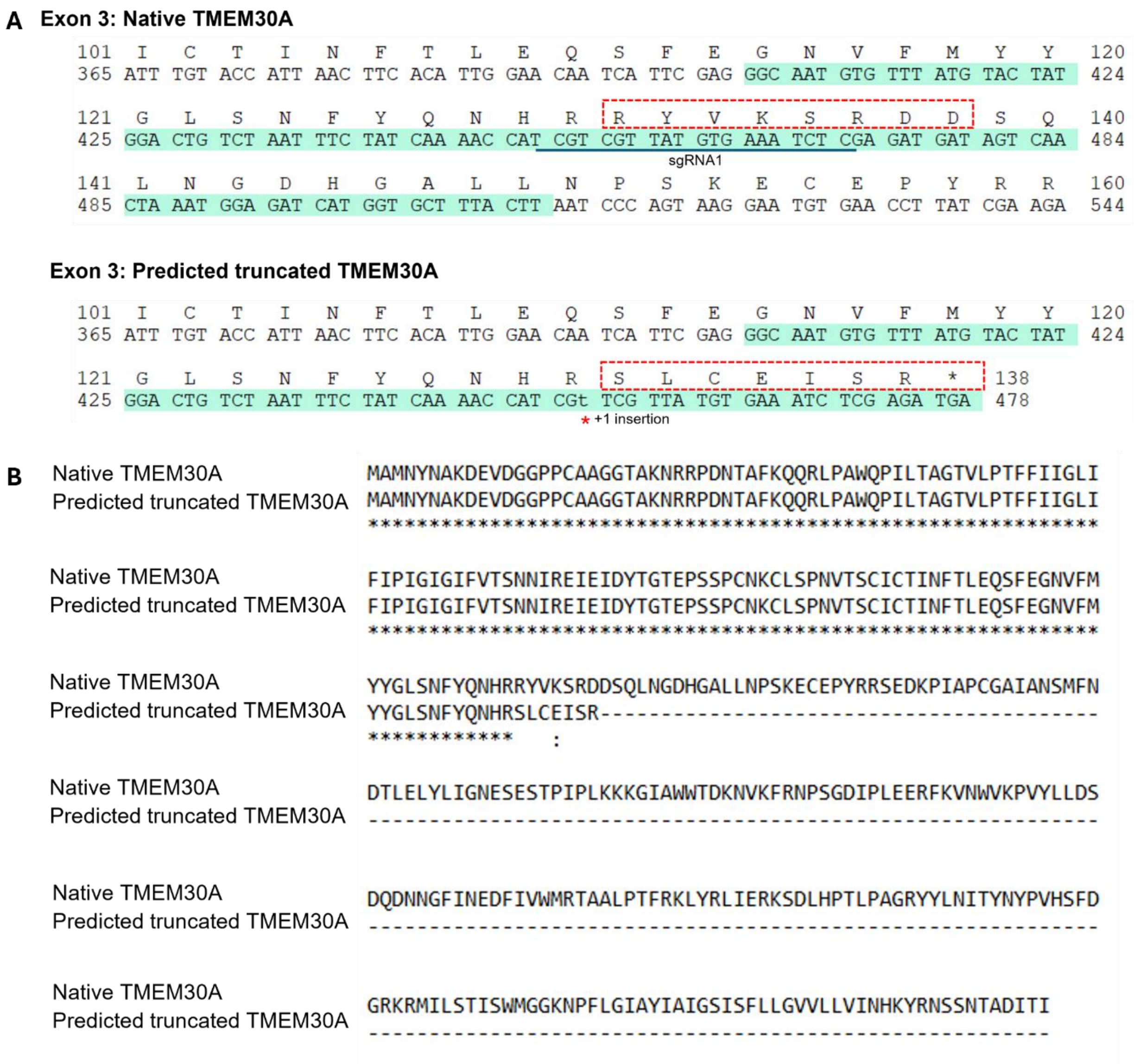


**Fig. S2** Analysis of predicted truncated TMEM30A protein sequence. (A) Nucleotide sequence of exon 3 is highlighted in green. The red boxes indicate the amino acids (aa) that are predicted to change following a single nucleotide insertion. (B) Comparison of protein sequences of native TMEM30A (356 aa) and predicted truncated TMEM30A (137 aa).
